# Ant nurse workers exhibit behavioral and transcriptomic signatures of specialization on larval stage

**DOI:** 10.1101/218834

**Authors:** Justin T. Walsh, Michael R. Warner, Adrian Kase, Benjamin J. Cushing, Timothy A. Linksvayer

## Abstract

Division of labor within and between the worker and queen castes is thought to underlie the tremendous success of social insects. Colonies might benefit if subsets of nurse workers specialize further in caring for larvae of a certain stage or caste, given that larval nutritional requirements depend on stage and caste. We used short- (<1 hr) and long-term (ten days) behavioral observations to determine whether nurses of the pharaoh ant (*Monomorium pharaonis*) exhibit such specialization. We found that nurses were behaviorally specialized based on larval instar but not on larval caste. This specialization was widespread, with 56% of nurses in the short-term and between 22-27% in the long-term showing significant specialization. Additionally, we identified ∼200 genes that were differentially expressed in nurse head and abdominal tissues between nurses feeding young versus old larvae. These included 18 genes predicted to code for secreted proteins, which may be passed from nurses to larvae via trophallaxis, as well as vitellogenin and major royal jelly protein-1, which have previously been implicated in the transfer of nutrition from nurse to larvae and the regulation of larval development and caste in social insects. Altogether, our results provide the first evidence in any social insect for a division of labor among nurse workers based on larval stage, and our study begins to elucidate the molecular mechanisms underlying this specialization.

## Introduction

Division of labor, one of the defining characteristics of eusociality, is believed to be the primary reason for the tremendous success of social insects (Wilson 1971; Oster and Wilson 1978; Wilson 1987). Within this system of division of labor, queens specialize on reproduction while workers specialize on tasks including brood care, foraging, and nest defense (Oster and Wilson 1978; Wilson 1987; Beshers and Fewell 2001). Increased worker efficiency within colonies is thought to be the main colony-level benefit of division of labor. Behavioral specialists, through learning or physiological differences, are expected to be more efficient than generalists (Oster and Wilson 1978; Robinson 1992; Wahl 2002), but see (Dornhaus 2008; Muscedere et al. 2009). Indeed, social insect behavioral specialists demonstrate increased efficiency in nest emigration (Langridge et al. 2008), nest excavation (Jeanson et al. 2008), undertaking (Trumbo and Robinson 1997; Julian and Cahan 1999), and response to sucrose (Perez et al. 2013).

Worker specialization is widespread, and is driven by a diversity of factors and proximate mechanisms. In many species, worker specialization depends on age, with younger workers generally performing tasks inside the nest (e.g. brood care) and older workers performing tasks outside the nest (e.g. foraging) (Oster and Wilson 1978; Robinson 1992; Beshers and Fewell 2001; Mikheyev and Linksvayer 2015). Alternatively, worker tasks can be allocated based on body size and shape, as many species exhibit morphologically distinct worker sub-castes that perform different roles within the colony (Oster and Wilson 1978; Beshers and Fewell 2001). Worker variation in behavioral specialization can also occur independently of age and morphology (Gordon 1989; Jeanson and Weidenmuller 2014). This interindividual variability can be the result of genetic diversity among workers (Oldroyd and Fewell 2007), environmental differences during early development (Tautz et al. 2003; Weidenmuller et al. 2009), variation in adult nutritional state (Blanchard et al. 2000; Ament et al. 2011; Charbonneau et al. 2017), prior experience (Theraulaz et al. 1998), and the social environment (Webster and Ward 2011).

Cooperative brood care, which includes feeding, grooming, and carrying brood, is one of the most important suites of tasks performed by adult workers (Oster and Wilson 1978; Wilson 1987). Different larvae have different nutritional requirements depending upon their caste and developmental stage (Cassill and Tschinkel 1996). For example, young larvae of many ant species are fed exclusively liquid food via nurse-larva trophallaxis while older larvae are also fed solid protein (Petralia et al. 1980; Tschinkel 1988; Cassill et al. 2005). Furthermore, old larvae require more frequent and longer feedings than young larvae (Cassill and Tschinkel 1996, 1999).

The caste fate of developing larvae in social insects is socially regulated by nurse workers (Linksvayer et al. 2011; Linksvayer 2015; Vojvodic et al. 2015), often based on the quantity and quality of nutrition provided to larvae (Wheeler 1986; Hunt and Nalepa 1994; Trible and Kronauer 2017). In ants, adult queens tend to have higher fat and protein content relative to workers, and it is usually assumed that queen-destined larvae are fed different quantities and qualities of food compared to worker-destined larvae (Hunt and Nalepa 1994; Smith and Suarez 2010; Amor et al. 2016; Warner et al. 2016). Furthermore, recent research in the Florida carpenter ant (*Camponotus floridanus)* found that nurse workers transfer juvenile hormone, microRNAs, hydrocarbons, various peptides, and other compounds to larvae during feeding (LeBoeuf et al. 2016), providing a potential further mechanism for nurses to provide stage- and caste-specific nutrition to larvae that may regulate larval development.

Recent research in honey bees (*Apis mellifera*) suggests that nurse workers exhibit both behavioral and transcriptomic specialization on larval caste (He et al. 2014; Vojvodic et al. 2015). However, these studies did not test for specialization on larval stage and, to the best of our knowledge, no previous study has investigated the potential for nurse specialization on caste or larval stage in ants. In this study, we tested whether individual pharaoh ant (*Monomorium pharaonis*) nurse workers exhibit behavioral specialization on different larval stages or castes, as measured on both short (< 1 hr) and long (10 days) timescales. We estimated how widespread such specialization is and the contribution of specialists to colony-level brood care. Building on our behavioral results, we used an existing transcriptomic data set to identify genes with expression patterns that may be associated with nurse specialization. Overall, we sought to elucidate whether nurse specialization exists in ants, how it contributes to colony-level brood care, and what gene expression patterns might be associated with such specialization.

## METHODS

### Background and Overall Design

All colonies used in this study were reared in the lab and were derived from stock colonies that have been systematically interbred for the past 10 years. We fed the colonies twice per week with an agar-based synthetic diet (Dussutour and Simpson 2008) and mealworms, and we maintained all colonies at 27 ± 1 °C, 50% relative humidity, and a 12:12 light:dark cycle. We conducted all behavioral observations manually using a dissecting microscope and red light. To keep the temperature constant during behavioral observations, we kept the colonies on a heating pad set to 27 °C.

*M. pharaonis* larvae have three instars (Alvares et al. 1993) that are distinguishable by body size, body shape, hair abundance, and hair morphology (Berndt and Kremer 1986). Although reproductive-destined larvae (males and gynes) cannot be distinguished from worker-destined larvae as eggs or 1st instar larvae, they can be readily distinguished after the 1st instar (Berndt and Kremer 1986; Edwards 1991). Since colonies usually only produce new gynes and males in the absence of fertile queens (Peacock et al. 1955; Edwards 1987), we set up queen-absent colonies, which rear both worker- and reproductive-destined larvae, when testing for specialization on larval caste. For both the behavioral observations and transcriptomic analyses, we initially classified the larvae into five stages based on size and hair morphology: 1st instar, 2nd instar, and small, medium, and large 3rd instar (see Berndt and Kremer 1986; Warner et al. 2016 for details). However, for subsequent behavioral analyses, we only considered larval instar.

### Short-term Observations

First we conducted short-term observations of unmarked workers in both queen-present (n = 8) and queen-absent (n = 3) colonies to determine whether nurses exhibited short-term specialization based on larval instar (using queen-present colonies) or larval caste (using queen-absent colonies). We observed colonies until we saw a worker feed a larva of any instar or caste, and then we continuously observed that nurse worker for as long as possible (max = 67 minutes). We recorded each time the nurse fed a larva, as well as the stage and caste of the larva, using the event logging software “BORIS” (Friard and Gamba 2016). We defined feeding behavior as a stereotypical behavioral interaction between the nurse worker and larva in which the mouthparts of the nurse and larva were in contact for at least three seconds. We defined both the transfer of solid food particles and liquid food via trophallaxis from nurse to larva as feeding behavior and did not distinguish between these two feeding behaviors. We restricted subsequent analysis to nurses we observed feeding at least times.

### Long-term Observations

Next, we attempted to test whether individually-marked nurses in queenless colonies express long-term specialization (across ten days). We wanted to track nurses for at least ten days because this time scale includes the entire amount of time that *M. pharaonis* workers tend to perform nursing behaviors (Mikheyev and Linksvayer 2015). In each of five colonies, we collected a cohort of 63 one-day-old callow workers and we uniquely painted each of these focal individuals with paint dots on their heads and abdomens using combinations of eight colors. Specifically, we lightly anesthetized them with carbon dioxide and marked their heads and abdomens with a dot of paint using Sharpie extra-fine point, oil-based paint pens (Dornhaus 2008; Dornhaus et al. 2008; Charbonneau et al. 2017). To control for potential behavioral effects of the paint, we painted all remaining adult workers in the colonies with black dots on their heads and abdomens. Because all 63 focal individuals in each colony were age-matched, we were able to control for possible effects of nurse age on potential behavioral specialization.

We constructed queen-absent colonies with 400 workers and 2.5 mL of brood (i.e. approximately 500 eggs, larvae, and pupae of different stages; Warner et al. 2016, 2018) and recorded all observed feeding, grooming, or carrying behaviors performed by all focal individually-marked workers. We initially used queen absent colonies because such colonies normally raise new queens and we wanted to test for longer-term specialization for caste. However, given that we observed no short-term specialization for caste, and our colonies ended up not producing sexual brood, we only considered potential long-term specialization based on larval stage. We defined feeding as described above, when an individually-marked worker’s mandibles interacted with a larva’s mandibles for at least three seconds. We defined grooming as an interaction between worker mandibles and a larva for a minimum of three seconds. We defined carrying as a worker lifting a larva with her mandibles and transporting the larva to another location. We analyzed feeding, grooming, and carrying behavior separately. We observed all colonies for three hours per day for ten consecutive days and restricted subsequent analysis to individuals we observed feeding, grooming, or carrying at least three times.

### Statistical Analysis of Behavioral Specialization

We performed all statistical analyses in R version 3.4.1 (R Core Team 2014). For both short and long-term observations, we first used binomial generalized linear mixed models (GLMMs) to ask whether individual nurses differed significantly in their degree of specialization on larval stage or caste. To test for nurse specialization on larval instar, we grouped 1st and 2nd instar larvae as “young” larvae and all 3rd instar larvae as “old” larvae. This grouping is biologically meaningful as 1st and 2nd instar larvae are fed solely a liquid diet while 3rd instar larvae are also fed solid food (Petralia et al. 1980; Tschinkel 1988; Cassill et al. 2005). Specifically, we fit GLMMs with the R package lme4 (Bates et al. 2015) for the proportion of fed larvae that were young versus old, with the identity of the nurse as a random effect and colony identity and nurse age as fixed effects when appropriate. Similarly, to test for nurse specialization on larval caste, we fit GLMMs for the proportion of fed larvae that were reproductive-versus worker-destined larvae. We evaluated the significance of both fixed and random effects using likelihood ratio (LR) tests. LR tests are appropriate for evaluating the significance of random effects in binomial models when the models contain fewer than three random effects (Bolker et al. 2009). A significant random effect of nurse identity in these models indicates that there is variation among individual nurses for degree of behavioral specialization, providing initial evidence for behavioral specialization within colonies.

Next, given that we found evidence for behavioral specialization (see Results), we used binomial tests to ask whether each individual significantly specialized on young versus old larvae, or reproductive versus worker larvae, based on recorded observations. We restricted analysis to nurses with at least six observations because this is the minimum number of observations that could potentially identify significant (P < 0.05) specialization with a binomial test. We estimated the expected frequency (i.e. “probability of success” in the binomial test) of interacting with larvae of one stage or caste relative to another stage or caste based on the observed proportion of interactions for the two stages or castes (e.g., the number of observed interactions between nurses and 1st instar larvae relative to 3rd instar larvae). In order to first determine whether any of the individual nurses we observed could be confidently classified as specialists, we first used binomial tests with a type I error rate corrected for multiple comparisons across all tested individuals. Given that some individuals were confidently identified as specialists with these conservative criteria, we next estimated the overall proportion of specialist versus non-specialist nurses in our study colonies using a type I error rate of 0.05 for each binomial test run separately for each individual nurse. This test provides an unbiased approach to determine one-at-a-time whether each individual displayed significant specialization or not.

### Gene Expression Analysis

Warner et al. (2017) performed RNA sequencing on a developmental time series of the five *M. pharaonis* larval stages as well as nurses collected in the act of feeding each of these larval stages. This previous study focused on identifying caste-biased genes across development and studying patterns of molecular evolution of these genes. In the current study, we take advantage of the fact that nurse samples used in Warner et al. (2017) were collected in the act of feeding one of the five larval stages, and we use the Warner et al. (2017) data set to compare transcriptomes of nurses feeding different larval stages. We chose to focus on nurses feeding very young versus very old larvae to maximize our power to detect differential expression based on the stage of larvae fed. Specifically, we used 11 samples of tissues from nurses collected in the act of feeding 1st instar larvae (5 head samples and 6 abdomen samples) and 10 samples of nurses collected in the act of feeding large 3rd instar larvae (5 head and 5 abdomen) to identify genes differentially expressed between nurses feeding larvae at the extreme young and old end of the developmental trajectory. See supplemental information for a brief summary of the sample collection procedure; for details of sample collection, RNA extraction, library preparation, sequencing, and estimation of per-locus expression, see Warner et al. (2017).

After removing lowly expressed genes (FPKM < 1 in ½ the samples), we used the package EdgeR (Robinson et al. 2010) for differential expression analysis. We constructed a GLM-like model, including larval stage fed, replicate, and queen presence as additive effects to identify genes differentially expressed between nurses feeding young versus old larvae (1st instar versus large 3rd instar; separately for head and abdomens). We calculated gene ontology (GO) term enrichment of differentially expressed genes using the R package GOstats, with a cut-off P-value of 0.05 (Falcon and Gentleman 2007).

To test whether genes found to be differentially expressed between nurses tended to code for secreted proteins in *Drosophila melanogaster*, we compiled a list of genes annotated as coding for secreted proteins according to the online tool GLAD (Hu et al. 2015). From this list, we identified secreted proteins with orthologs in *M. pharaonis* using a recently created orthology map between *M. pharaonis*, *Apis mellifera*, and *D. melanogaster* (Warner et al. unpublished manuscript; orthology map included as Supplementary Data). We estimated the association between a gene’s likelihood to be differentially expressed and secreted, removing all genes for which a *D. melanogaster* ortholog was not detected. We generated plots using the R package ggplot2 (Wickham 2009).

## RESULTS

### Short-term Specialization on Larval Stage

We observed 52 nurses feed at least three times (mean = 8.8 feeding events) and we included these nurses in the GLMMs. The random effect of nurse identity was significant, suggesting that nurses tended to specialize on feeding either young or old larvae (table 1). Next, to classify each individual nurse as showing significant specialization or not (i.e. to classify nurses as specialists or non-specialists), we used binomial tests with an expected proportion of old larvae relative to young plus old larvae of 0.781 (the observed proportion of old larvae fed across all individuals in long-term feeding observations). We used the observed proportion from long-term observations, as opposed to short-term observations, because we specifically attempted to balance the number of recorded old and young larvae feeding events (in terms of total number of observations, not per individual) during short-term observations. Therefore the observed short-term proportions are not an accurate representation of the naturally-occurring proportions. We included the 32 nurses we observed feed at least six times. When using a type I error rate corrected for multiple comparisons, which should produce a conservative estimate of the frequency of specialists across the whole study, we classified about 56% (18/32) of nurses as specialists (Bonferroni-adjusted P). When using a type I error rate of 0.05, which should yield an unbiased estimate of the frequency of specialists versus generalists within colonies, we again classified about 56% (18/32) of nurses as specialists. These specialists performed about 65% (242/375) of the observed feedings.

**Table 1.** Summary of effects of factors on short- and long-term nurse behavior on likelihood ratio tests of GLMMs

| | $\chi^2$ | df | p |
| --- | --- | --- | --- |
| Short-term Feeding |  |  |  |
| Caste |  |  |  |
| Individual Nurse | 1.430 | 1 | 0.232 |
| Stage |  |  |  |
| Individual Nurse | 345.56 | 1 | <0.0001 |
| Long-term Feeding |  |  |  |
| Individual Nurse | 36.934 | 1 | <0.0001 |
| Colony | 57.750 | 4 | <0.0001 |
| Age | 0.018 | 1 | 0.892 |
| Long-term Grooming |  |  |  |
| Individual Nurse | 21.357 | 1 | <0.0001 |
| Colony | 49.576 | 1 | <0.0001 |
| Age | 0.022 | 1 | 0.882 |
| Long-term Carrying |  |  |  |
| Individual Nurse | 4.500 | 1 | 0.034 |
| Colony | 5.048 | 1 | 0.025 |
| Age | 0.021 | 1 | 0.886 |

### Short-term Specialization on Caste

We observed 22 nurses feed at least three times (mean = 5.64 feeding events). The random effect of nurse identity in the GLMM was not significant (table 1) indicating that nurses did not specialize on larval caste. In the binomial tests, we included the ten nurses we observed feed at least six times and used an expected proportion of reproductive-destined larvae of 0.534. When correcting for multiple comparisons, we classified zero nurses as specialists. When using a type I error rate of 0.05, we classified 10% (1/10) of nurses as specialists and this specialist performed about 6% (9/142) of the observed feedings (figure 1).

**Figure 1.**
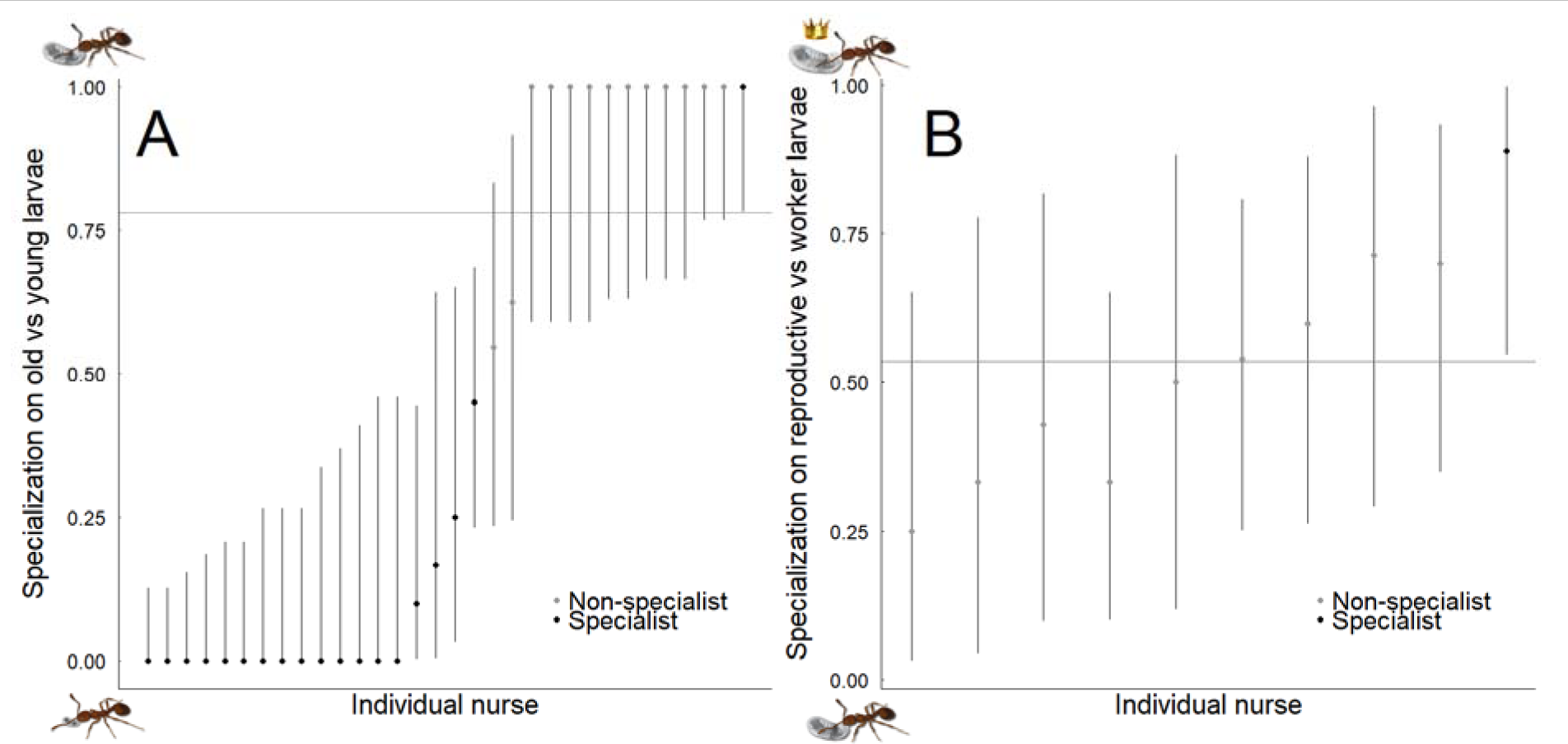
Short-term nurse worker specialization on young vs old larvae (A) and worker- vs reproductive-destined larvae (B). The dots represent the proportions of old larvae (A) or reproductive larvae (B) that each nurse worker fed and the error bars are the confidence intervals from the binomial tests. The horizontal line represents the expected proportion based on overall observed proportion of interactions. In plot A, a proportion of 1 means the nurse worker fed only old larvae while a 0 means the worker fed only young larvae. In plot B, a proportion of 1 means the nurse worker fed only reproductive-destined larvae while a 0 means the worker fed only worker-destined larvae.

### Long-term Feeding Specialization on Larval Stage

We observed 40 nurses feed at least three times (mean = 12.9 feeding events). The effects of nurse identity and colony identity were significant (table 1), indicating that nurses tended to specialize on feeding either young or old larvae. The age of the nurse was not significant. In the binomial tests, we included the 30 nurses we observed feed at least six times and used an expected proportion of old larvae of 0.781. When correcting for multiple comparisons, we classified 20% (6/30) of nurses as being long-term specialists on larval stage. When using an uncorrected type I error rate of 0.05, we classified about 27% (8/30) of nurses as long-term specialists and these long-term specialists performed about 42% (201/480) of the observed feedings (figure 2). Long-term specialists performed significantly more feedings than non-specialists (Mann-Whitney-Wilcoxon test; W= 19.5, P = 0.0013).

**Figure 2.**
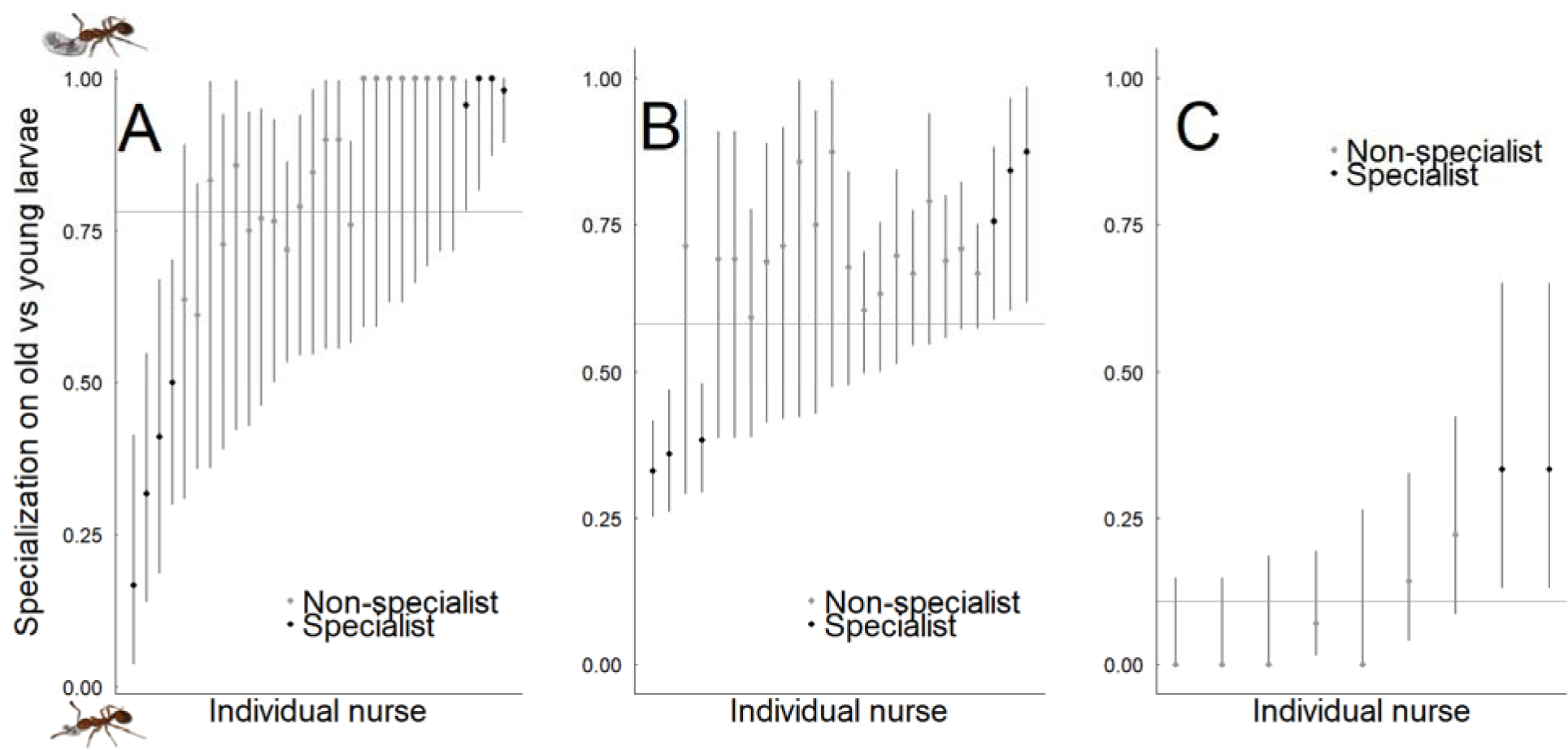
Nurse worker specialization on young vs old larvae for long-term feeding (A), grooming (B), and carrying (C). The dots represent the proportions of old larvae that each nurse worker cared for and the error bars are the confidence intervals from the binomial tests. The horizontal line represents the expected proportion based on overall observed proportion of interactions. A proportion of 1.0 means the nurse worker cared for only old larvae while a 0 means the worker cared for only young larvae.

### Long-term Grooming Specialization on Larval Stage

We observed 32 individuals grooming larvae at least three times (mean = 33.9 grooming events). The effects of nurse identity and colony identity were significant (table 1), indicating that nurses tended to specialize on feeding either young or old larvae. The age of the nurse was not significant. In the binomial tests, we included the 24 nurses we observed groom at least six times and used an expected proportion of old larvae of 0.581. When correcting for multiple comparisons, we classified about 13% (3/24) of nurses as specialists. When using an uncorrected type I error rate of 0.05, we classified 25% (6/24) of nurses as specialists and these specialists performed about 39% (406/1053) of the observed groomings (figure 2). The number of groomings performed by specialists and non-specialists was not significantly different (W = 29, P = 0.1021).

### Long-term Carrying Specialization on Larval Stage

We observed 17 individuals carrying a larva at least three times (mean = 13.4 carrying observations). The effects of nurse identity and colony identity were significant (table 1), indicating that nurses tended to specialize on feeding either young or old larvae. The age of nurse was not significant. In the binomial tests, we included the nine nurses we observed carrying a larva at least six times and used an expected ratio of old to young of 0.107. When correcting for multiple comparisons, we classified zero nurses as specialists. When using an uncorrected type I error rate of 0.05, we classified about 22% (2/9) of nurses as specialists and these specialists performed about 12% (24/197) of the carrying observations (figure 2). The number of groomings performed by specialists and non-specialists was not significantly different (W = 13, P = 0.100).

### Transcriptomic Analysis

We identified 209 and 173 differentially expressed genes (DEGs) in the heads and abdomens, respectively, of nurses collected while feeding young (i.e. 1st instar) versus old (i.e. large 3rd instar) worker larvae (FDR < 0.05; figure 3*a,b*). In both nurse heads and abdomens, we identified more up-regulated genes in nurses feeding young versus old larvae (two-sided binomial, null hypothesis of 50% upregulated in nurses feeding young larvae; heads: N = 209, P < 0.001; abdomens: N = 173, P < 0.001). There was a positive association between genes up-regulated in the heads and genes upregulated in the abdomens of nurses feeding young larvae (χ^2^ = 312, df = 1, P < 0.001), as well as between genes up-regulated in the heads and abdomens of nurses feeding old larvae (χ^2^ = 260, df = 1, P < 0.001). Additionally, there was an overall correlation between expression fold change in nurse heads and abdomens across all differentially expressed genes between nurses feeding young versus old larvae (figure 3*c*). For genes associated with each nurse type, gene ontology was largely dominated by metabolism-related categories (table A1). Genes up-regulated in the heads of nurses feeding young larvae were also associated with isoprenoid (a type of hydrocarbon) processing, and genes up-regulated in the abdomens of nurses feeding young larvae were associated with transport and localization.

**Figure 3.**
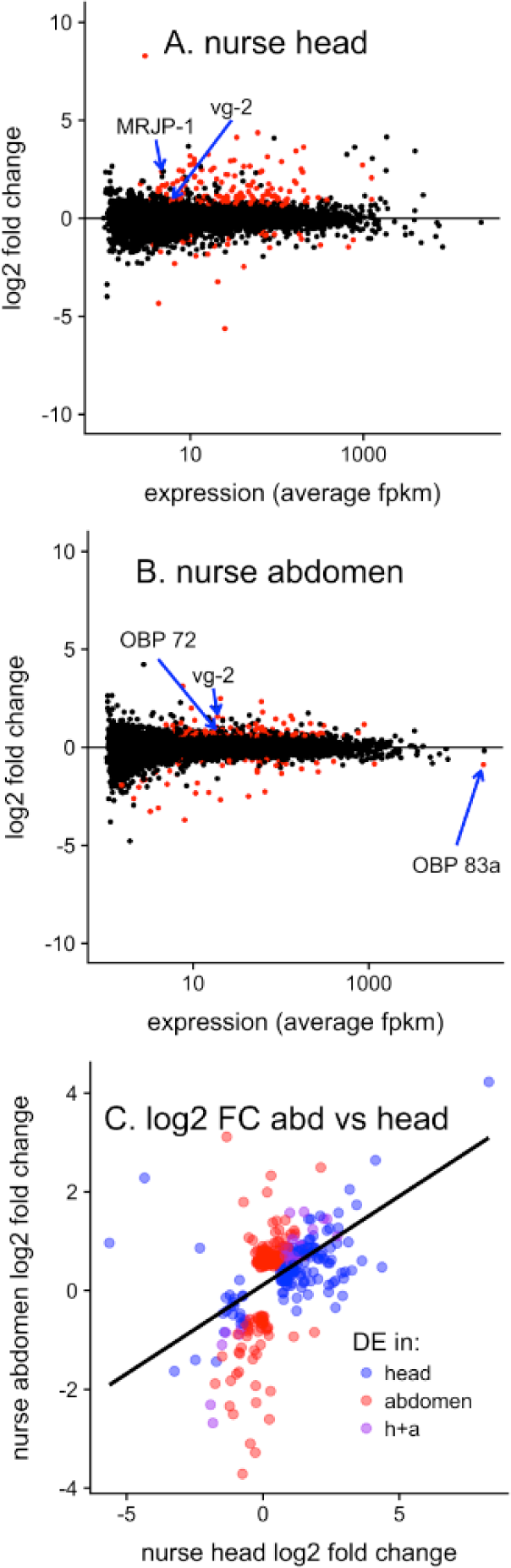
Differential expression between nurses feeding young (1st-instar) and old (large 3rd instar) larvae in A) nurse heads and B) nurse abdomens. Genes colored red are differentially expressed (FDR < 0.05). C) Correlation of log2 fold change of differentially expressed genes as measured in nurse abdomens and heads (Spearman rho = 0.345, P < 0.001). Black line represents trendline of linear model. Genes are colored by tissue differentially expressed in (FDR < 0.05). In all plots, genes with positive “log2 fold change” are upregulated in nurses feeding large 1st vs 3rd instar larvae (i.e feeding young vs old larvae).

Genes that were differentially expressed in nurses based on larval stage were more likely to code for proteins known to be cellularly secreted in *Drosophila melanogaster* (χ^2^ = 29.1, df = 1, P < 0.001; 18 DGEs coding for secreted proteins out of 148 total DGEs with orthologs in *D. melanogaster*. 178 genes had orthologs that code for secreted proteins in *D. melanogaster*, out of 5391 genes in the analysis). Nearly all of the DEGs that are predicted to code for secreted proteins were upregulated in nurses feeding young larvae (14/14 in heads, 9/10 in abdomens, see table A2 for complete list of DEGs based on larval stage fed).

## DISCUSSION

The tremendous ecological success of social insects is thought to be primarily due to efficient division of labor within colonies (Wilson 1971; Oster and Wilson 1978; Wilson 1987). Here we provide to the best of our knowledge the first evidence for the existence of a division of labor within nurse workers based on the instar of larvae they care for. We found evidence for behavioral specialization in the short-term (less than an hour) and the long-term (over 10 days). Nurses specialized on either old (3rd instar) or young (1st and 2nd instar) larvae, and this specialization was consistent across feeding, grooming, and carrying behaviors. In the short-term, we classified 56% of nurses as specialists in terms of feeding, and in the long-term, we classified 27%, 25%, and 22% of workers as specialists in feeding, grooming, and carrying respectively.

Specialists are predicted to increase colony efficiency (Oster and Wilson 1978; Robinson 1992; Wahl 2002). Although we cannot say whether specialist nurses increase *M*. *pharaonis* colony efficiency, our data suggest that specialists do play an important role in the colony as they performed more per capita feedings than non-specialists. The specialization of nurse workers on old or young larvae might be explained by specialization on trophallaxis (i.e. feeding liquids) versus feeding solid food particles since young larvae are fed only a liquid diet while old larvae are also fed solid protein (Petralia et al. 1980; Tschinkel 1988; Cassill et al. 2005). If so, the nurses specialized for trophallaxis may play a disproportionately large role in regulating larval development since trophallactic fluid contains not only nutrition but also juvenile hormone, microRNAs, hydrocarbons, various peptides, and other compounds (LeBoeuf et al. 2016).

Given that we found evidence for the behavioral specialization of nurses on young versus old larvae, we also tested for differential gene expression in the head and abdominal tissues of nurses feeding young versus old larvae as a first step in identifying transcriptomic signatures of specialization. We expect that differentially expressed genes (DEGs) might be functionally associated with different types of care provided by nurse workers to differently-aged larvae, which might actively contribute to the social regulation of larval development (see Vojvodic et al. 2015). We identified 209 and 173 differentially expressed genes in nurse heads and abdomens, respectively, between nurses feeding young versus old worker-destined larvae. Based on our behavioral analyses, we expect approximately one half of the individuals used in our gene expression samples to be specialized based on larval age, rendering our differential expression analysis effectively conservative (i.e. non-specialists included in our sample would weaken the transcriptomic signature of specialists). Interestingly, while the majority of DEGs were tissue specific, a significant proportion of genes were differentially expressed in the same direction in heads and abdomens, and there was an overall correlation between log fold expression change from young to old nurses in both heads and abdomens. Together, these results indicate that some transcriptomic changes associated with nurse specialization occur throughout nurse bodies.

Intriguingly, genes with *D. melanogaster* orthologs that are known to code for cellularly-secreted proteins were overrepresented among the DEGs between *M. pharaonis* nurses feeding young versus old larvae. The DEGs we detected in nurse tissues could directly affect larval development if the proteins were secreted by nurses and transferred to larvae via trophallaxis (Linksvayer 2015). Many of these DEGs which are predicted to code for secreted proteins have metabolic functions, suggesting they may play a role in the the breakdown of food before it is passed to larvae.

The DEGs between nurses may also play a role in responding to larval signals. Two odorant binding proteins (OBP) were differentially expressed in nurse abdomens (figure 3*a*). These OBPs potentially play a role in communication between nurses and larvae (Zhou et al. 2015, McKenzie et al. 2016). Although OBPs are predicted to be primarily expressed in the antennae, previous studies found that OBPs are frequently expressed in non-chemosensory tissues (McKenzie et al. 2014) and can exhibit various functions beyond olfaction (Nomura et al. 1992, Maleszka et al. 2007, Dani et al. 2011, Zhang et al. 2016). For example, the Gp-9 gene encodes for the odorant binding protein SiOBP3 and has been linked to colony organization in the fire ant *Solenopsis invicta* (Wang et al. 2013). Expression of SiOBP3 is found throughout the bodies of workers, gynes, and males and is actually lowest in the antennae (Zhang et al. 2016).

Interestingly, in both measured tissues (heads and abdomens), we identified more genes upregulated in nurses feeding 1st-instar larvae than those feeding 3rd-instar larvae, and all DEGs that code for proteins secreted in *D. melanogaster* were upregulated in nurses feeding 1st-instar larvae. These genes might be involved in regulating early larval development, or perhaps regulation of larval caste fate, given that caste determination occurs at least by the end of the 1st instar (Berndt and Kremer 1986; Alvares et al. 1993; Khila et al. 2010; Warner et al. 2016; Warner et al. 2018). Genes upregulated in nurses feeding 1st-instar larvae included genes such as vitellogenin (Vg2) (Libbrecht et al. 2013) and a member of the major royal jelly protein family (MRJP-1) (Schonleben et al. 2007), both of which have been implicated in the production and transfer of proteinaceous food to honey bee larvae, which then shapes larval development and caste fate (Amdam et al. 2003) (figure 3*a*). Interestingly, LeBoeuf et al (2016) found both a MRJP homolog and vitellogenin in the trophallactic fluid of ant nurses fed to developing larvae. Therefore, it is possible that *M. pharaonis* nurses feeding young larvae are passing on these compounds directly to larvae as a means to regulate larval development.

Contrary to findings in honey bees (He et al. 2014; Vojvodic et al. 2015), we found no behavioral evidence for nurse specialization on larval caste. This lack of specialization in *M. pharaonis* is somewhat surprising, given that worker- and reproductive-destined larvae likely have different nutritional needs (Hunt and Nalepa 1994; Smith and Suarez 2010; Amor et al. 2016; Warner et al. 2016). However, this difference may be attributable to differences in timing of caste determination. In honey bees, caste determination occurs relatively late in development and over a period of time, as queen-worker inter-castes can be produced by experimental manipulation of diet late in development (Linksvayer et al. 2011; Dedej et al. 1998; Wang et al. 2014). Therefore, in honey bees, continued nurse-larval interactions are likely essential to fine-tune caste dimorphism (Linksvayer et al. 2011).

In contrast to honey bees where each larva develops in an isolated brood cell, worker- and reproductive-destined larvae are not spatially separated in *M. pharaonis*. This lack of separation could also help explain the lack of specialization on larval caste in *M. pharaonis* compared to honey bees. Additionally, many ants (including *M. pharaonis*) spatially arrange their brood such that younger larvae and eggs tend be in the center and older larvae and pupae towards the edge of brood piles (Franks & Sendova-Franks 1992; Lim & Lee 2005). This spatial arrangement could potentially contribute to the observed short-term specialization if nurses spend most of their time in one area of the nest and feed larvae close to them. However, we observed nurses frequently moving around the nest during our short-term observations, interacting with other workers or collecting food in between subsequent feedings, so that each individual nurse had the potential to interact with all brood stages.

Further research is necessary to characterize the implications of nurse specialization, elucidate the detailed molecular and physiological underpinnings, and to determine how widespread specialization is across ants and other social insects. Interestingly, we found significant effects of the colony identity for long-term nursing, grooming, and carrying. Although outside the scope of this study, it is possible that different colonies exhibit different levels of specialization in either the number of specialists or the proportion of brood care behaviors performed by specialists. Future studies should test for colony-level variation in nurse specialists.

### Conclusion

This study describes a previously undocumented form of division of labor within ant nurse workers: specialization based on larval instar. We found evidence for this specialization in three different brood care behaviors. Additionally, we found ∼200 differentially expressed genes between nurses feeding young versus old larvae. Contrary to findings in honey bees, we found no evidence for specialization of nurse workers on larval caste. Further research is necessary to characterize the implications of nurse specialization, elucidate the detailed molecular and physiological underpinnings, and to determine how widespread specialization is across ants and other social insects.

### Data Accessibility

Data supporting this paper are included as supplemental files.

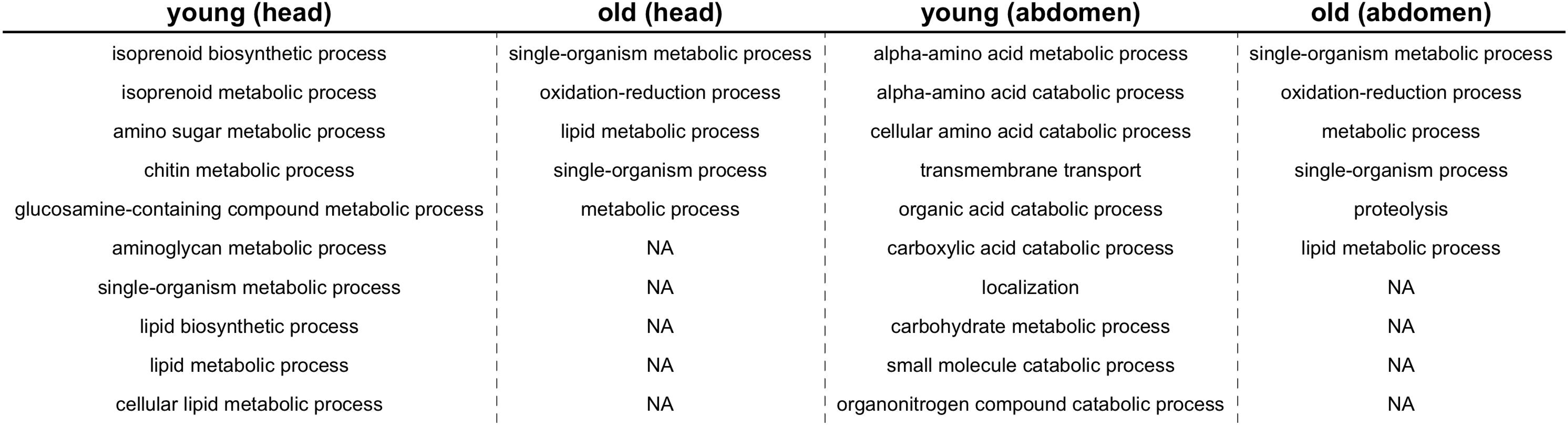

## REFERENCES

Alvares, L. E., Bueno, O. C., & Fowler, H. G. (1993). Larval instars and immature development of a Brazilian population of pharaohs ant, *Monomorium pharaonis*. [Article]. Journal of Applied Entomology-Zeitschrift Fur Angewandte Entomologie, 116(1), 90–93.

Amdam, G. V., Norberg, K., Hagen, A., & Omholt, S. W. (2003). Social exploitation of vitellogenin. [Article]. Proceedings of the National Academy of Sciences of the United States of America, 100(4), 1799–1802. doi: 10.1073/pnas.0333979100

Ament, S. A., Chan, Q. W., Wheeler, M. M., Nixon, S. E., Johnson, S. P., Rodriguez-Zas, S. L., … Robinson, G. E. (2011). Mechanisms of stable lipid loss in a social insect. [Article]. Journal of Experimental Biology, 214(22), 3808–3821. doi: 10.1242/jeb.060244

Amor, F., Villalta, I., Doums, C., Angulo, E., Caut, S., Castro, S., … Boulay, R. (2016). Nutritional versus genetic correlates of caste differentiation in a desert ant. [Article]. Ecological Entomology, 41(6), 660–667. doi: 10.1111/een.12337

Bates, D., Machler, M., Bolker, B. M., & Walker, S. C. (2015). Fitting Linear Mixed-Effects Models Using lme4. [Article]. Journal of Statistical Software, 67(1), 1–48.

Berndt, K. P., & Kremer, G. (1986). Larvenmorphologie der Pharoameise *Monomorium pharaonis*. Zoologischer Anzeiger.

Beshers, S. N., & Fewell, J. H. (2001). Models of division of labor in social insects. [Review]. Annual Review of Entomology, 46, 413–440. doi: 10.1146/annurev.ento.46.1.413

Blanchard, G. B., Orledge, G. M., Reynolds, S. E., & Franks, N. R. (2000). Division of labour and seasonality in the ant *Leptothorax albipennis*: worker corpulence and its influence on behaviour. [Article]. Animal Behaviour, 59, 723–738. doi: 10.1006/anbe.1999.1374

Bolker, B. M., Brooks, M. E., Clark, C. J., Geange, S. W., Poulsen, J. R., Stevens, M. H. H., & White, J. S. S. (2009). Generalized linear mixed models: a practical guide for ecology and evolution. [Review]. Trends in Ecology & Evolution, 24(3), 127–135. doi: 10.1016/j.tree.2008.10.008

Cassill, D. L., Butler, J., Vinson, S. B., & Wheeler, D. E. (2005). Cooperation during prey digestion between workers and larvae in the ant, *Pheidole spadonia*. [Article]. Insectes Sociaux, 52(4), 339–343. doi: 10.1007/s00040-005-0817-x

Cassill, D. L., & Tschinkel, W. R. (1996). A duration constant for worker-to-larva trophallaxis in fire ants. [Article]. Insectes Sociaux, 43(2), 149–166. doi: 10.1007/bf01242567

Cassill, D. L., & Tschinkel, W. R. (1999). Regulation of diet in the fire ant, *Solenopsis invicta*. [Article]. Journal of Insect Behavior, 12(3), 307–328.

Charbonneau, D., Poff, C., Nguyen, H., Shin, M. C., Kierstead, K., & Dornhaus, A. (2017). Who are the “lazy” ants? The function of inactivity in social insects and a possible role of constraint: inactive ants are corpulent and may be young and/or selfish. Integrative and Comparative Biology, 57(3), 649–667.

Dani, F. R., Michelucci, E., Francese, S., Mastrobuoni, G., Cappellozza, S., La Marca, G., … Pelosi, P. (2011). Odorant-Binding Proteins and Chemosensory Proteins in Pheromone Detection and Release in the Silkmoth *Bombyx mori*. [Article]. Chemical Senses, 36(4), 335–344. doi: 10.1093/chemse/bjq137

Dedej, S., Hartfelder, K., Aumeier, P., Rosenkranz, P., & Engels, W. (1998). Caste determination is a sequential process: effect of larval age at grafting on ovariole number, hind leg size and cephalic volatiles in the honey bee (*Apis mellifera* carnica). [Article]. Journal of Apicultural Research, 37(3), 183–190.

Dornhaus, A. (2008). Specialization does not predict individual efficiency in an ant. PLoS Biology, 6(11).

Dornhaus, A., Holley, J. A., Pook, V. G., Worswick, G., & Franks, N. R. (2008). Why do not all workers work? Colony size and workload during emigrations in the ant *Temnothorax albipennis*. [Article]. Behavioral Ecology and Sociobiology, 63(1), 43–51. doi: 10.1007/s00265-008-0634-0

Dussutour, A., & Simpson, S. J. (2008). Description of a simple synthetic diet for studying nutritional responses in ants. [Article]. Insectes Sociaux, 55(3), 329–333. doi: 10.1007/s00040-008-1008-3

Edwards, J. P. (1987). Caste regulatin in the pharoahs ant *Monomorium pharaonis*-The influence of queens on the production of new sexual forms. [Article]. Physiological Entomology, 12(1), 31–39. doi: 10.1111/j.1365-3032.1987.tb00721.x

Edwards, J. P. (1991). Caste regulation in the pharaohs ant *Monomorium pharaonis*-recognition and cannibalism of sexual brood by workers. Physiological Entomology, 16(3), 263–271.

Falcon, S., & Gentleman, R. (2007). Using GOstats to test gene lists for GO term association. [Article]. Bioinformatics, 23(2), 257–258. doi: 10.1093/bioinformatics/btl567

Franks, N. R., & Sendova-Franks, A. B. (1992). Brood sorting by ants- distributing the workload over the work-surface. [Article]. Behavioral Ecology and Sociobiology, 30(2), 109–123.

Friard, O., & Gamba, M. (2016). BORIS: a free, versatile open-source event-logging software for video/audio coding and live observations. [Article]. Methods in Ecology and Evolution, 7(11), 1325–1330. doi: 10.1111/2041-210x.12584

Gordon, D. M. (1989). Dynamics of task switching in harvester ants. [Article]. Animal Behaviour, 38, 194–204. doi: 10.1016/s0003-3472(89)80082-x

He, X. J., Tian, L. Q., Barron, A. B., Guan, C., Liu, H., Wu, X. B., & Zeng, Z. J. (2014). Behavior and molecular physiology of nurses of worker and queen larvae in honey bees (*Apis mellifera*). [Article]. Journal of Asia-Pacific Entomology, 17(4), 911–916. doi: 10.1016/j.aspen.2014.10.006

Hu, Y., Comjean, A., Perkins, L. A., Perrimon, N., & Mohr, S. E. (2015). GLAD: an Online Database of Gene List Annotation for *Drosophila*. J Genomics, 3, 75–81. doi: 10.7150/jgen.12863

Hunt, J. H., & Nalepa, C. A. (1994). Nourishment, evolution, and insect sociality. Westview, Boulder: Westview, Boulder.

Jeanson, R., Clark, R. M., Holbrook, C. T., Bertram, S. M., Fewell, J. H., & Kukuk, P. F. (2008). Division of labour and socially induced changes in response thresholds in associations of solitary halictine bees. [Article]. Animal Behaviour, 76, 593–602. doi: 10.1016/j.anbehav.2008.04.007

Jeanson, R., & Weidenmuller, A. (2014). Interindividual variability in social insects - proximate causes and ultimate consequences. [Review]. Biological Reviews, 89(3), 671–687. doi: 10.1111/brv.12074

Julian, G. E., & Cahan, S. (1999). Undertaking specialization in the desert leaf-cutter ant *Acromyrmex versicolor*. [Article]. Animal Behaviour, 58, 437–442. doi: 10.1006/anbe.1999.1184

Khila, A., & Abouheif, E. (2010). Evaluating the role of reproductive constraints in ant social evolution. [Article]. Philosophical Transactions of the Royal Society B-Biological Sciences, 365(1540), 617–630. doi: 10.1098/rstb.2009.0257

Langridge, E. A., Sendova-Franks, A. B., & Franks, N. R. (2008). How experienced individuals contribute to an improvement in collective performance in ants. [Article]. Behavioral Ecology and Sociobiology, 62(3), 447–456. doi: 10.1007/s00265-007-0472-5

LeBoeuf, A. C., Waridel, P., Brent, C. S., Goncalves, A. N., Menin, L., Ortiz, D., … Keller, L. (2016). Oral transfer of chemical cues, growth proteins and hormones in social insects. [Article]. Elife, 5, 27. doi: 10.7554/eLife.20375

Libbrecht, R., Corona, M., Wende, F., Azevedo, D. O., Serrao, J. E., & Keller, L. (2013). Interplay between insulin signaling, juvenile hormone, and vitellogenin regulates maternal effects on polyphenism in ants. [Article]. Proceedings of the National Academy of Sciences of the United States of America, 110(27), 11050–11055. doi: 10.1073/pnas.1221781110

Lim, S. P., & Lee, C. Y. (2005). Brood arrangement and food distribution among larvae under different colony conditions in the Pharaoh’s ant, *Monomorium pharaonis* (Hymenoptera: Formicidae). [Article]. Sociobiology, 46(3), 491–503.

Linksvayer, T. A. (2015). The Molecular and Evolutionary Genetic Implications of Being Truly Social for the Social Insects. In A. Zayed & C. F. Kent (Eds.), Genomics, Physiology and Behaviour of Social Insects (Vol. 48, pp. 271–292). London: Academic Press Ltd-Elsevier Science Ltd.

Linksvayer, T. A., Kaftanoglu, O., Akyol, E., Blatch, S., Amdam, G. V., & Page, R. E. (2011). Larval and nurse worker control of developmental plasticity and the evolution of honey bee queen-worker dimorphism. [Article]. Journal of Evolutionary Biology, 24(9), 1939–1948. doi: 10.1111/j.1420-9101.2011.02331.x

Maleszka, J., Foret, S., Saint, R., & Maleszka, R. (2007). RNAi-induced phenotypes suggest a novel role for a chemosensory protein CSP5 in the development of embryonic integument in the honeybee (*Apis mellifera*). [Article]. Development Genes and Evolution, 217(3), 189–196. doi: 10.1007/s00427-006-0127-y

McKenzie, S. K., Fetter-Pruneda, I., Ruta, V., & Kronauer, D. J. C. (2016). Transcriptomics and neuroanatomy of the clonal raider ant implicate an expanded clade of odorant receptors in chemical communication. [Article]. Proceedings of the National Academy of Sciences of the United States of America, 113(49), 14091–14096. doi: 10.1073/pnas.1610800113

McKenzie, S. K., Oxley, P. R., & Kronauer, D. J. C. (2014). Comparative genomics and transcriptomics in ants provide new insights into the evolution and function of odorant binding and chemosensory proteins. [Article]. Bmc Genomics, 15, 14. doi: 10.1186/1471-2164-15-718

Mikheyev, A. S., & Linksvayer, T. A. (2015). Genes associated with ant social behavior show distinct transcriptional and evolutionary patterns. [Article]. Elife, 4, 29. doi: 10.7554/eLife.04775

Muscedere, M. L., Willey, T. A., & Traniello, J. F. A. (2009). Age and task efficience in the ant *Pheidole dentata*: young minor workers are not specialist nurses. Animal Behaviour, 77, 911–918.

Nomura, A., Kawasaki, K., Kubo, T., & Natori, S. (1992). Purification and localization of P10, a novel protein that increases in nymphal regenerating legs of *Periplaneta americana* (American cockroach). [Article]. International Journal of Developmental Biology, 36(3), 391–398.

Oldroyd, B. P., & Fewell, J. H. (2007). Genetic diversity promotes homeostasis in insect colonies. [Review]. Trends in Ecology & Evolution, 22(8), 408–413. doi: 10.1016/j.tree.2007.06.001

Oster, G. F., & Wilson, E. O. (1978). Caste and ecology in the social insects. Princeton, NJ: Princeton University Press.

Peacock, A. D., Sudd, J. H., & Baxter, A. T. (1955). Studies in pharaohs ant, *Monomorium pharaonis*. Entomologist’s Monthly Magazine, 91, 130–133.

Perez, M., Rolland, U., Giurfa, M., & d’Ettorre, P. (2013). Sucrose responsiveness, learning success, and task specialization in ants. [Article]. Learning & Memory, 20(8), 417–420. doi: 10.1101/lm.031427.113

Petralia, R. S., Sorensen, A. A., & Vinson, S. B. (1980). Labial gland system of the larvae of the imported fire ant, *Solenopsis invicta* Buren- Ultrastructure and enzyme analysis [Article]. Cell and Tissue Research, 206(1), 145–156.

Robinson, G. E. (1992). Regulation of division-of-labor in insect societies. [Review]. Annual Review of Entomology, 37, 637–665. doi: 10.1146/annurev.en.37.010192.003225

Robinson, M. D., McCarthy, D. J., & Smyth, G. K. (2010). edgeR: a Bioconductor package for differential expression analysis of digital gene expression data. [Article]. Bioinformatics, 26(1), 139–140. doi: 10.1093/bioinformatics/btp616

Schonleben, S., Sickmann, A., Mueller, M. J., & Reinders, J. (2007). Proteome analysis of Apis mellifera royal jelly. [Article]. Analytical and Bioanalytical Chemistry, 389(4), 1087–1093. doi: 10.1007/s00216-007-1498-2

Smith, C. R., & Suarez, A. V. (2010). The trophic ecology of castes in harvester ant colonies. [Article]. Functional Ecology, 24(1), 122–130. doi: 10.1111/j.1365-2435.2009.01604.x

Tautz, J., Maier, S., Groh, C., Rossler, W., & Brockmann, A. (2003). Behavioral performance in adult honey bees is influenced by the temperature experienced during their pupal development. [Article]. Proceedings of the National Academy of Sciences of the United States of America, 100(12), 7343–7347. doi: 10.1073/pnas.1232346100

R Core Team. 2014 R: a language and environment for statistical computing. R Foundation for Statistical Computing, Vienna, Austria. http://www.R-project.org/.

Theraulaz, G., Bonabeau, E., & Deneubourg, J. L. (1998). Response threshold reinforcement and division of labour in insect societies. [Article]. Proceedings of the Royal Society B- Biological Sciences, 265(1393), 327–332.

Trible, W., & Kronauer, D. J. C. (2017). Caste development and evolution in ants: it’s all about size. Journal of Experimental Biology, 220(1), 53–62. doi: 10.1242/jeb.145292

Trumbo, S. T., & Robinson, G. E. (1997). Learning and task interference by corpse-removal specialists in honey bee colonies. [Article]. Ethology, 103(11), 966–975.

Tschinkel, W. R. (1988). Social control of egg laying rate in queens of the fire ant, *Solenopsis invicta*. [Article]. Physiological Entomology, 13(3), 327–350. doi: 10.1111/j.1365- 3032.1988.tb00484.x

Vojvodic, S., Johnson, B. R., Harpur, B. A., Kent, C. F., Zayed, A., Anderson, K. E., & Linksvayer, T. A. (2015). The transcriptomic and evolutionary signature of social interactions regulating honey bee caste development. Ecology and Evolution, 5(21), 4795–4807. doi: 10.1002/ece3.1720

Wahl, L. M. (2002). Evolving the division of labour: Generalists, specialists and task allocation. [Article]. Journal of Theoretical Biology, 219(3), 371–388. doi: 10.1006/jtbi.2002.3133

Wang, J., Wurm, Y., Nipitwattanaphon, M., Riba-Grognuz, O., Huang, Y. C., Shoemaker, D., & Keller, L. (2013). A Y-like social chromosome causes alternative colony organization in fire ants. [Article]. Nature, 493(7434), 664–668. doi: 10.1038/nature11832

Wang, Y., Kaftanoglu, O., Fondrk, M. K., & Page, R. E. (2014). Nurse bee behaviour manipulates worker honeybee *(Apis mellifera* L.) reproductive development. [Article]. Animal Behaviour, 92, 253–261. doi: 10.1016/j.anbehav.2014.02.012

Warner, M. R., Kovaka, K., & Linksvayer, T. A. (2016). Late-instar ant worker larvae play a prominent role in colony-level caste regulation. [Article]. Insectes Sociaux, 63(4), 575–583. doi: 10.1007/s00040-016-0501-3

Warner, M. R., Lipponen, J., & Linksvayer, T. A. (2018). Pharaoh ant colonies dynamically regulate reproductive allocation based on colony demography. [journal article]. Behavioral Ecology and Sociobiology, 72(3), 31. doi: 10.1007/s00265-017-2430-1

Warner, M. R., Mikheyev, A. S., & Linksvayer, T. A. (2017). Genomic signature of kin selection in an ant with obligately sterile workers. [Article]. Molecular Biology and Evolution, 34(7), 1780–1787. doi: 10.1093/molbev/msx123

Webster, M. M., & Ward, A. J. W. (2011). Personality and social context. [Article]. Biological Reviews, 86(4), 759–773. doi: 10.1111/j.1469-185X.2010.00169.x

Weidenmuller, A., Mayr, C., Kleineidam, C. J., & Roces, F. (2009). Preimaginal and adult experience modulates the thermal response behavior of ants. [Article]. Current Biology, 19(22), 1897–1902. doi: 10.1016/j.cub.2009.08.059

Wheeler, D. E. (1986). Developmental and physiological determinants of caste in social hymenoptera- Evolutionary implications. [Article]. American Naturalist, 128(1), 13–34. doi: 10.1086/284536

Wickham, H. (2009). ggplot2: Elegant graphics for data analysis. New York, New York: Spring-Verlag.

Wilson, E. O. (1971). The insect societies. Cambridge, Massachusetts: Harvard University Press.

Wilson, E. O. (1987). Causes of ecological success- the case of the ants. [Article]. Journal of Animal Ecology, 56(1), 1–9. doi: 10.2307/4795

Zhang, W., Wanchoo, A., Ortiz-Urquiza, A., Xia, Y. X., & Keyhani, N. O. (2016). Tissue, developmental, and caste-specific expression of odorant binding proteins in a eusocial insect, the red imported fire ant, *Solenopsis invicta*. [Article]. Scientific Reports, 6, 16. doi: 10.1038/srep35452

Zhou, X. F., Rokas, A., Berger, S. L., Liebig, J., Ray, A., & Zwiebel, L. J. (2015). Chemoreceptor Evolution in Hymenoptera and Its Implications for the Evolution of Eusociality. [Article]. Genome Biology and Evolution, 7(8), 2407–2416. doi: 10.1093/gbe/evv149

